# Separable roles for RanGTP in nuclear and ciliary trafficking of a kinesin-2 subunit

**DOI:** 10.1101/562272

**Authors:** Shengping Huang, Prachee Avasthi

## Abstract

Kinesin is part of the microtubule (MT)-binding motor protein superfamily, which exerts crucial functions in cell division and intracellular transport in different organelles. The heterotrimeric kinesin-2, consisting of the kinesin like protein KIF3A/3B heterodimer and kinesin-associated protein KAP3, is highly conserved across species from the green alga *Chlamydomonas* to humans. It plays diverse roles in cargo transport including anterograde (base to tip) trafficking in cilium. However, the molecular determinants mediating trafficking of heterotrimeric kinesin-2 itself is poorly understood. Using the unicellular eukaryote *Chlamydomonas* and mammalian cells, we show that RanGTP regulates ciliary trafficking of KAP3. We found the armadillo repeat region 6-9 (ARM6-9) of KAP3, required for its nuclear translocation, is sufficient for its targeting to the ciliary base. Given that KAP3 is essential for cilia formation and the emerging roles of RanGTP/importin β in ciliary protein targeting, we further investigate the effect of RanGTP in cilium length regulation in these two different systems. We demonstrate that precise control of RanGTP levels, revealed by different Ran mutants, is crucial for cilium formation and maintenance. Most importantly, we were able to segregate RanGTP regulation of ciliary protein incorporation from of its nuclear roles. Our work provides important support for the model that nuclear import mechanisms have been coopted for independent roles in ciliary import.

## Introduction

Cilia are microtubule-based protrusions with sensory and/or motile functions. In mammals, defects in assembly and maintenance of cilia results in a series of diseases called “ciliopathies” (Fliegauf et al., 2007; Mitchison et al., 2017; Anvarian et al., 2019; Breslow et al.,2019). The assembly and maintenance of these organelles are dependent on anterograde and retrograde intraflagellar transport (IFT) (Rosenbaum et al., 2002). Anterograde IFT, which moves from the base of a cilium to the tip, is driven by kinesin-2 (kozminski et al., 1995; Cole et al., 1998), whereas retrograde IFT, which moves from the tip back to the base, is achieved by cytoplasmic dynein 1b (Signor et al., 1999; Pazour et al., 1999; Porter et al., 1999). The kinesin-2 motor family is composed of a heterotrimeric KIF3A/KIF3A/KAP3 motor and a homodimeric KIF17 motor (Hirokawa et al. 2009). Unlike homodimeric KIF17, heterotrimeric kinesin-2 is highly conserved, and loss of function in any component of heterotrimeric kinesin-2 results in defective cilia in different organisms (Walther et al., 1994; Morris et al., 1997; Sarpal et al., 2004; Zhao et al., 2011). In addition to cilium formation, regulation and maintenance of cilium length is also dependent on the size and frequency of kinesin-2 trains recruited to/entering cilia (Ludington et al., 2013; Engel et al., 2009). In addition to its central role in IFT and ciliogenesis, heterotrimeric kinesin-2 has also been reported in other organelle transport events outside cilia. This includes anterograde transport from endoplasmic reticulum to the Golgi apparatus in *Xenopus* (Le Bot et al., 1998), retrograde transport from the Golgi to endoplasmic reticulum in HeLa cells (Stauber et al., 2006), and establishment of cell polarity during migration (Murawala et al., 2009). Furthermore, heterotrimeric kinesin-2 is reported to play critical roles in mitosis (Fan et al., 2004; Haraguchi et al., 2006).

Cilia assemble in quiescent cells and disassemble in dividing cells (Plotnikova et al., 2009). In quiescent cells, heterotrimeric kinesin-2 is localized in both cilia and basal body in ciliated cells. In dividing cells, when cells enter mitosis and cilia retract, the non-motor subunit KAP is transported into nucleus before nuclear membrane break down in cells of sea urchin blastulae (Morris et al., 2004). During cytokinesis, the motor protein KIF3B is localized in the midbody (Fan et al., 2004). Macromolecules can’t freely go into or out of the cilium and nucleus because diffusion barriers exist at the ciliary base and the nuclear pore complex (NPC) (Kee et al., 2012; Breslow et al., 2013; Takao et al. 2014., Endicott et al., 2018). The fundamental question is how this conserved heterotrimeric kinesin-2 complex traffics between different compartments (the cytoplasm, nucleus and cilium) for specific functions, and how these processes are regulated.

For nuclear transport from the cytoplasm, the NPC mediates active transport of proteins (Alber et al., 2007; Wente SR et al., 2010; Strambio-De-Castillia et al., 2010; Beck et al., 2017). Larger proteins (>50 kDa) generally require RanGTP and specific importin transport receptors to cross the NPC (Gorlich et al., 1994; Gorlich and vogel et al., 1995). Importins usually bind a nuclear localization signal (NLS)-containing cargo at relatively low RanGTP level in cytosol. This moves the complex through NPCs and releases cargo in the nucleus where RanGTP concentration is high (Moore et al.,1993; Gorlich et al., 1999). In most cases, importin β1 binds to importin α, which interacts with a conventional NLS, to mediate substrate nuclear import (Gorlich and Kostka et al., 1995). In contrast, importin β2 recognizes non-traditional proline-tyrosine NLS (PY-NLS) for nuclear import (Lee et al., 2006). In the nucleus, direct binding of RanGTP with importin β results in cargo release (Gorlich et al., 1997).

Compared to nucleo-cytoplasmic transport, the molecular mechanisms or sequence motifs controlling membrane or soluble protein trafficking into cilia are less well understood. Previous studies reported that several cis-acting elements, including RVxP, VxPx, and Ax[S/A]xQ motifs, are important for mediating ciliary trafficking of membrane proteins (Jenkins et al., 2006; Geng et al., 2006; Mazelova et al., 2009; Berbari et al., 2008). Several recent studies implicated that the cilium and nucleus are co-evolved for signal integration because of the shared components and evolutionary origin (Mcclure-Begley et al., 2017; Johnson et al., 2019; Satir et al., 2019). It was also proposed that there are some shared mechanisms between ciliary import and nuclear import (Dishinger et al., 2010; Takao et al., 2014; Del Viso et al., 2016; Takao et al., 2017; Endicott et al., 2018). Several results indicated that RanGTP/importin β/NLS import system is required for ciliary targeting of either membrane or soluble proteins including importin β1 for Crumsb3 (Fan et al., 2007), RanGTP/importin β2 for KIF17 and Gli2 (Tarrado et al., 2016; Dishinger et al., 2010), importin β2 for RP2 (Hurd et al., 2011), importin α1/α6/NLS for KIF17 (Funabashi et al., 2017), and importin β2/PY-NLS for GLI2/GLI3 (Han et al., 2017). It was also reported that importin β2/Rab8 forms a ternary complex with ciliary localization sequences to direct membrane protein trafficking to cilia (Madugula et al., 2016), suggesting that this process is independent of RanGTP and a NLS or PY-NLS. Despite these advancements in uncovering mechanisms for ciliary import, it remains unclear whether RanGTP regulates ciliary trafficking directly or by affecting nuclear import to result in defective ciliary trafficking. It is also unclear whether different cargoes depend on different importin β receptors for ciliary trafficking. Lastly, prior work investigated determinants of ciliary entry for the homodimeric ciliary kinesin composed of KIF17 (Dishinger et al., 2010), the heterotrimeric kinesin-2 is the motor that dictates cilium assembly and maintenance (Walther et al., 1994; Morris et al., 1997; Sarpal et al., 2004; Zhao et al., 2011; Engel et al., 2009; Ludington et al., 2013). Therefore, we focused on how ciliary trafficking of the heterotrimeric kinesin-2 is regulated by leveraging the unique advantages of the unicellular green alga *Chlamydomonas reinhardtii* as an excellent eukaryotic model to study ciliogenesis (Harris et al., 2001; Rosenbaum et al., 2002).

Cilia of *Chlamydomonas* can be regenerated to full length in two hours, and unlike mammalian cells, ciliary assembly does not need to be induced (Rosenbaum et al., 1969) to result in heterogeneous population of ciliated and non-ciliated cells. The molecular mechanism of the nucleo-cytoplasmic trafficking is likely conserved between the *Chlamydomonas* and humans (Li et al., 2018). However, there are fewer constituent nucleoporins in the *Chlamydomonas* NPC compared to that of humans (Neumann et al., 2006). By using both mammalian and *Chlamydomonas* cells, we have found that ciliary trafficking of kinesin associated protein KAP3 is regulated by RanGTP. We demonstrated that precise manipulation of RanGTP level is crucial for regulating cilium formation. Importantly, we were able to clearly show that RanGTP plays a direct role in incorporation of ciliary proteins that is independent of its nuclear roles. These results provide potential insights for the molecular mechanism orchestrating multi-compartment trafficking of the heterotrimeric kinesin-2 motor complex. Further, they answer a long-standing open question in the field about whether nuclear import mechanisms have been coopted for direct ciliary import.

## Results

### Ciliary protein KAP3 can localize to the nucleus

The heterotrimeric kinesin-2 motor complex consists of the heterodimeric motor proteins KIF3A/3B and the adaptor protein KAP3. In contrast, the homodimeric kinesin-2 KIF17 motor does not need an adaptor protein to exert its function. Although it was suggested that KAP3 functions as a linker between KIF3A/3B and the specific cargoes to facilitate intracellular transport, the function of KAP3 is still not well characterized. To explore this, we firstly investigated the intracellular localization of KAP3 in ciliated and non-ciliated cells. HA-tagged KAP3A and KAP3B (a short isoform of KAP3A) were transfected into hTERT-RPE cells. 24 hours after transfection, cilia were induced by serum starvation. As expected, both HA-tagged KAP3A and KAP3B co-localized with acetylated-α-tubulin, confirming the intracellular localization of KAP3A and KAP3B are not affected by small HA epitope and can be targeted to cilia (**Figure 1A**). Surprisingly, we noticed that a significant amount of KAP3A and KAP3B was also distributed throughout the nucleus (**Figure 1A**). We further examined the localization of KAP3A and KAP3B in other different cell types. As shown in **Figure 1B**, HA-tagged KAP3A and KAP3B are mainly localized in the nucleus of COS-7 cells, although a small amount is distributed in the cytoplasm. We also showed that EGFP-tagged KAP3A and KAP3B could localize in the nucleus of MDCK cells **(Figure 1C)**. Nuclear localization of KAP3A was consistent with that of EGFP tagged KAP3A in a previous report (Tenny et al., 2016).

**Figure 1.**
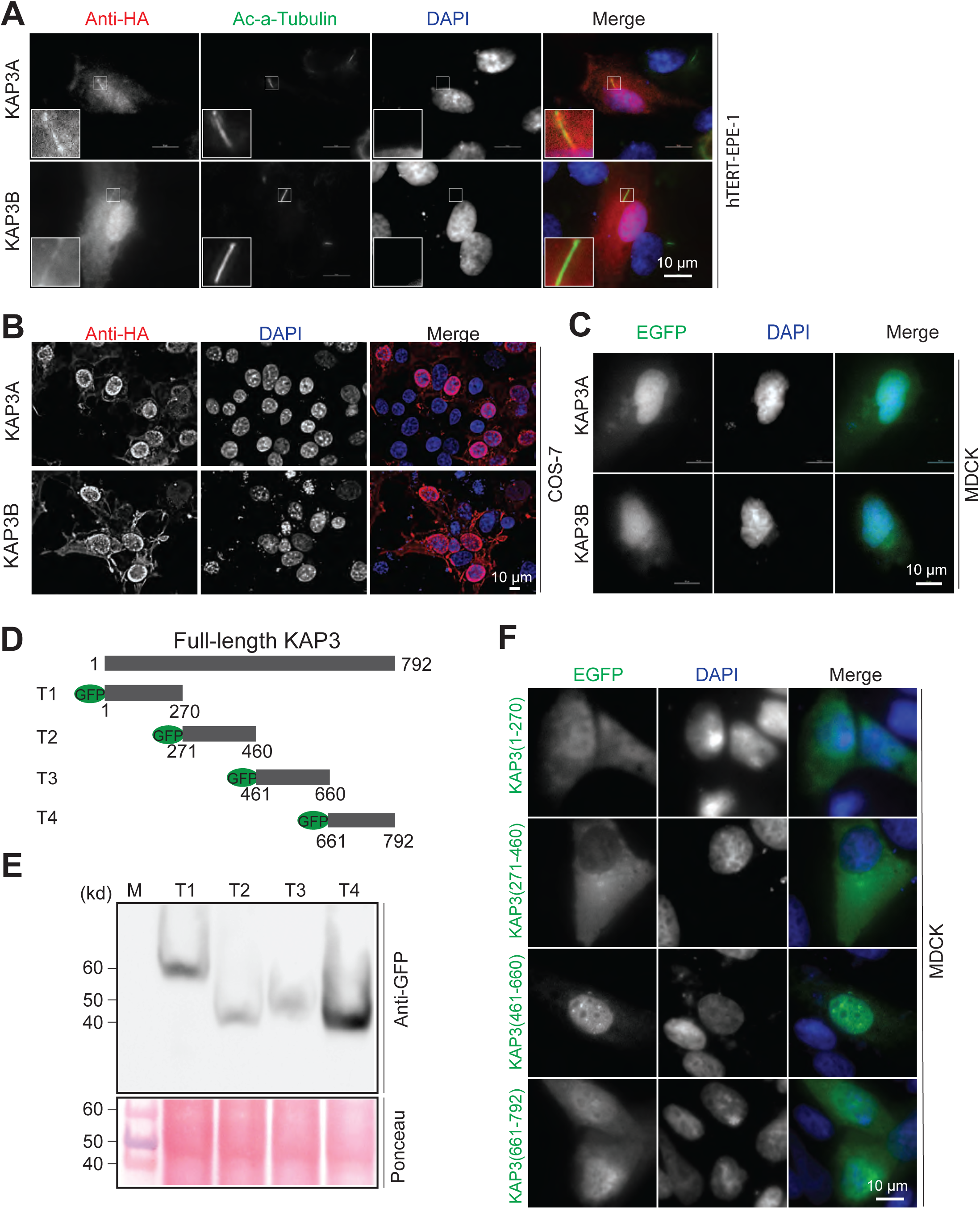
Kinesin-associated protein KAP3 localizes in both the cilium and nucleus. **A.** HA-tagged KAP3A or KAP3B were transfected into hTERT-RPE cells. After 24 hours transfection, cilium was induced under 0.25% serum starvation for another 24 hours. Cells were fixed with 4% paraformaldehyde (PFA) and stained with rabbit anti-HA and mouse anti-ac-α-tubulin antibodies**. B.** HA-tagged KAP3A or KAP3B were expressed in non-ciliated COS-7 cells, After 24 hour transfection, cells were fixed with 4% PFA and stained with anti-HA antibody. Cell nuclei are pseudo-colored blue, following staining with DAPI. **C**. EGFP tagged KAP3A or KAP3B were expressed in MDCK cells. **D.** Schematic illustration of EGFP tagged KAP3 truncated derivatives. **E**. Western blotting analysis of the expression of EGFP tagged KAP3 transiently transfected MDCK cells. **F**. Mapping the domains required for nuclear localization of KAP3A in MDCK cells: a series of EGFP tagged KAP3A truncations were transfected into MDCK cells. 24 hours after transfection, cells were fixed with 4% PFA and stained with mouse anti-HA antibody and DAPI.

To dissect critical regions within KAP3A responsible for its nuclear localization, as depicted in **Figure 1D**, EGFP-fused KAP3 truncations were constructed (henceforth, KAP3 refers to the long isoform KAP3A). First, the expression of appropriately-sized truncations in MDCK cells was detected by western blot analysis (**Figure 1E**). Second, the subcellular distributions of these KAP3 truncations in MDCK cells were analyzed via fluorescence microscopy. As shown in **Figure 1F**, the N-terminal fragment KAP3 (1-270) and the C-terminal fragment KAP3 (661-792) are distributed in both the cytoplasm and nucleus, and KAP3 (271-460) was exclusively distributed in the cytoplasm. In contrast, KAP3 (461-660), consisting of armadillo repeats (ARM) 6-9, were predominantly localized in the nucleus, which is similar to full-length KAP3. These data indicate that the region between amino acids 461 and 660 is crucial for nuclear localization of KAP3. Taken together, our data demonstrate that ciliary protein KAP3 can localize to the nucleus under the control of armadillo repeats 6-9.

### RanGTP, but not importin β2, mediates nuclear translocation of KAP3

As shown in **Figure 1**, KAP3 is distributed to the nucleus in different cells. To determine the molecular mechanism of KAP3 nuclear translocation, we tested a well-studied pathway for protein nuclear import, RanGTP mediated nuclear import, which requires a high concentration of RanGTP in the nucleus for the disassembly of the imported complexes. To determine whether RanGTP drives nuclear import of KAP3, the dominant negative mutant RanQ69L which cannot hydrolyze GTP, was used in this study. As shown in **Figure 2A**, Ectopic expression of RanQ69L blocked nuclear localization of KAP3 in COS-7 cells, resulting in a more cytoplasmic distribution of KAP3 relative to wild-type controls. This data suggests that nuclear translocation of KAP3 is mediated by a RanGTP-dependent nuclear import pathway.

**Figure 2.**
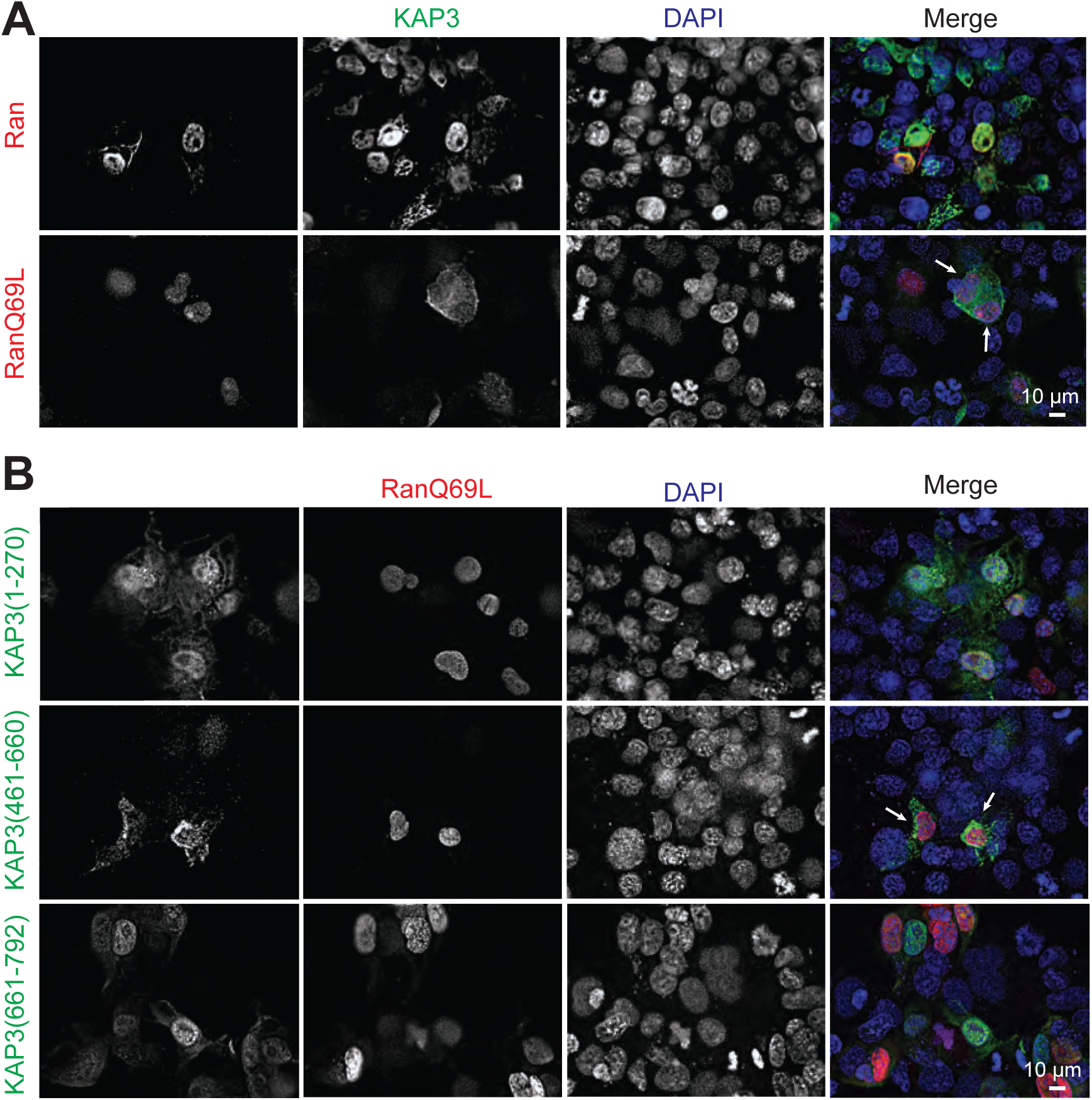
RanGTP regulates nuclear translocation of KAP3. **A.** Dominant negative form of Ran blocked nuclear localization of KAP3 in COS-7 cells. mCherry tagged wild-type Ran or GTP bound dominant mutant RanQ69L were co-transfected with EGFP-KAP3 into COS-7 cells. After 24 h transfection, cells were fixed with 4% PFA and stained with anti-EGFP antibody and DAPI **B.** Mapping the domains within KAP3 that is mediated by RanGTP mediated nuclear import. a series of EGFP tagged KAP3 truncations were co-transfected with GTP-locked Ran mutant Q69L into COS-7 cells. After 24 h transfection, cells were fixed with 4% PFA and stained with anti-EGFP antibody and DAPI.

We mapped the region responsible for nuclear localization of KAP3 and found that KAP3 (461-660) is required. If the region we mapped is correct, the nuclear localization of this truncation KAP3 (461-660) should be RanGTP-dependent, which act the same way as full-length KAP3. To test this, we co-transfected KAP3 (461-660) with the dominant negative Ran mutant RanQ69L. As shown in **Figure 2B**, RanQ69L completely disrupted nuclear import of KAP3 (461-660) and resulted in the cytoplasmic localization of this truncation. These results suggest that RanGTP-dependent nuclear import of KAP3 is dependent on the 461-660 region of KAP3. In contrast, RanQ69L didn’t change the localization of other KAP3 truncations (**Figure 2B**).

It was reported that the import receptor importin β2 plays critical roles in both nuclear import and ciliary import of ciliary proteins, like KIF17 and GLI2/GLI3 (Dishinger et al., 2010; Han et al., 2017). We further examined whether importin β2 is utilized for nuclear translocation of KAP3. The importin β2 inhibitory peptide M9M was used in these studies (Cansizoglu et al., 2007). hnRNP A1 was used as a positive control for this assay. As shown in **Figure S1A**, expression of MBP-tagged M9M inhibitory peptide disrupted nuclear localization of hnRNPA1 (*white arrow*), which confirmed that hnRNP A1 utilizes transport receptor importin β2 for its nuclear translocation. Compared to the empty MBP control, MBP-tagged M9M did not block the nuclear translocation of KAP3 **(Figure S1B).** This data suggests nuclear import of KAP3 is independent of the importin β2 receptor.

### The armadillo repeat domain 6-9 (ARM6-9) alone is sufficient for ciliary base localization

KAP3 can localize in both the cilium and nucleus. We dissected the regions required for nuclear translocation of KAP3. Next, we mapped the regions required for ciliary targeting of KAP3 in hTERT-RPE cells. As depicted in **Figure 3A**, full-length KAP3 contains three regions: a non-conserved N-terminal domain, nine armadillo repeats, and a C-terminal conserved domain (Jimbo et al., 2002; Shimuzu et al., 1996). Based on this, a series of truncations of KAP3 were generated and intracellular localization of these truncations was examined after cilium induction. As shown in **Figure 3B**, the truncation KAP3 (661-792) containing the C-terminal domain completely abolished localization to the cilium and ciliary base. In contrast, the truncation KAP3(186-792) containing both the nine armadillo repeats and C-terminal domain, and the truncation KAP3(186-660) merely with the nine armadillo repeats showed intense signal at the ciliary base. These data suggest that the nine armadillo repeats are required for KAP3 targeting to the ciliary base. We further narrowed the region within the nine armadillo repeats and demonstrated that the truncation KAP3(461-660), harboring the ARM6-9, is sufficient for ciliary base targeting of KAP3 (**Figure 3B**). It is noteworthy that this region is also required for RanGTP mediated KAP3 nuclear trafficking.

**Figure 3.**
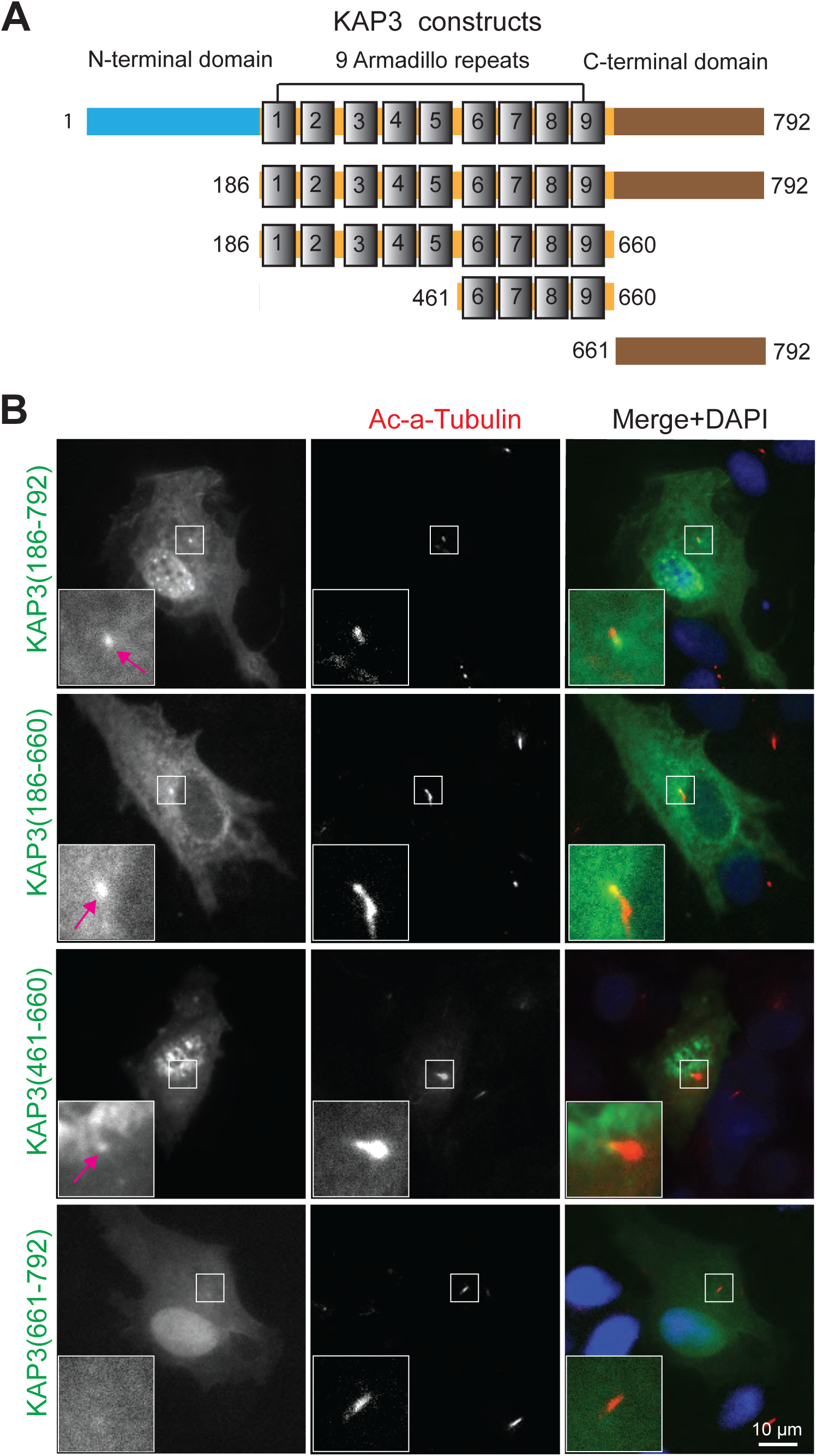
Armadillo repeat domain 6-9 targets KAP3 to the ciliary base. **A.** Schematic illustration of full-length human KAP3 and its truncated derivatives for analyzing ciliary targeting. **B.** Ciliary base localization of KAP3 truncations in hTERT-RPE cells. hTERT-RPE cells were transfected with various truncated constructs of KAP3. After 24 h transfection, the cells suffered serum starvation for cilium induction for another 24 hours and fixed with 4% PFA and co-stained with mouse anti-acetylated α-tubulin (red) and rabbit anti-EGFP antibodies (green). nuclei are stained with DAPI (blue).

### RanGTP regulates percent ciliation in human retinal epithelial cells

Given the middle region of KAP3, 461-660, is required for both nuclear and cilium base targeting and nuclear targeting is RanGTP dependent, we wanted to investigate whether KAP3-dependent ciliogenesis and cilium length regulation (Sarpal et al., 2003; Mueller et al., 2004), was also RanGTP dependent. We were further interested in Ran-dependent ciliary phenotypes and KAP3 localization due to previously reported shared mechanisms between nuclear and ciliary import processes (Dishinger et al., 2010; Takao et al., 2014; Del Viso et al., 2016; Takao et al., 2017; Endicott et al., 2018) and conflicting conclusions about the effect of RanGTP on ciliogenesis (Dishinger et al., 2010; Fan et al., 2011; Torrado et al., 2016). Wild-type Ran and three well-characterized dominant negative Ran mutants (RanQ69L, RanG19V and RanT24N) were used in this study. First, we studied the intracellular localization of these proteins in hTERT-RPE cells in serum-starved condition which induce ciliogenesis. As shown in **Figure S2**, all the mutants are predominantly localized in the nucleus which is similar to that of wild-type Ran. These data indicate that expression of these point mutants did not dramatically affect intracellular localization of Ran. To analyze the role of these mutants in cilium formation and length regulation in hTERT-RPE cells, wild type and mutant Ran expression plasmids were transfected into hTERT-RPE cells. 24 hours post-transfection, low serum media were added for 24 hours to induce cilium formation. As shown in **Figure 4A** and **4C**, ectopic expression of either the GTP locked mutants RanQ69L/RanG19V or the GDP-locked mutant RanT24N had no obvious effect on cilium length. In contrast, cells transfected with these dominant negative mutants could reduce ciliation percentage compared the un-transfected control cells **(Figure 4B)**. Further, different Ran mutants had different effects on ciliation percentage. RanQ69L has higher affinity to GTP and resulted in a dramatically reduction in ciliation percentage compared to RanG19V, which has relatively low affinity to GTP (Lounsburg et al., 1996) (**Figure 4B**). These data indicate that the RanGTP level in hTERT-RPE cells is a determinant of initiation of cilium formation. Taken together, the ability to bind and hydrolyze GTP by Ran in vivo, revealed by different dominant Ran mutants, regulates its essential functions on the generation of cilia.

**Figure 4.**
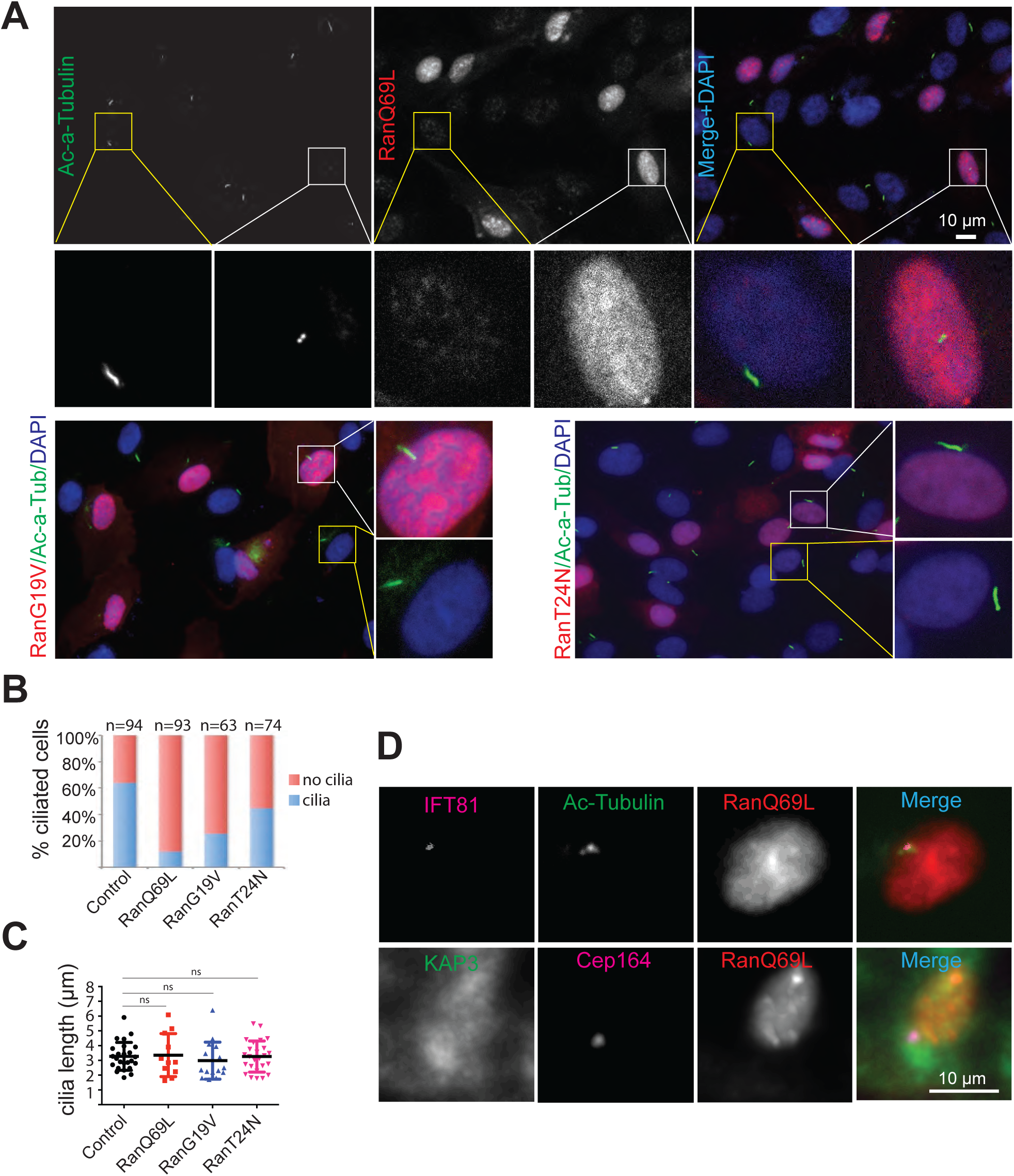
Dominant negative Ran mutants reduced percent ciliation in hTERT-RPE cells. **A.** dominant negative Ran mutants blocked cilia formation in hTERT-RPE cells. The plasmids expressing Ran mutant were transfected into hTERT-RPE cells. 24 h after transfection, Cilium was induced via serum starvation for another 24 h. The cells were fixed and labeled for acetylated α-tubulin and nuclei are stained with DAPI. **B.** hTERT-RPE cells expressing dominant Ran mutant reduced the percentage of ciliated cells. **C**. Quantification of cilia length. Data are presented as the mean±SEM. The unpaired *t*-test analysis was performed. *P*-value of great than 0.05 was considered no significant difference. The experiment was repeated three times. **D**. Localization of IFT81 and KAP3 in ciliated hTERT-RPE cells expressing RanQ69L. Cilium was induced in hTERT-RPE cells expressing RanQ69L via serum starvation. The cells were fixed and co-stained with acetylated α-tubulin and IFT81, or KAP3 and Cep164. Nuclei are stained with DAPI.

To determine why there is reduced cilium formation in RanQ69L-expressing RPE cells, we further investigated the localization of other important components, like the IFT complex and kinesin-2. As shown in **Figure 4D**, IFT81, a component of the IFT-B complex, is still localized in the ciliary base of hTERT-RPE cells expressing RanQ69L, suggesting that IFT-B targeting is not affected and is unlikely to be the primary cause of defective cilium formation. In contrast, the kinesin-2 associated protein KAP3 didn’t localize to the ciliary base. This data suggests that, in addition to potential roles for RanGTP in ciliary entry, ciliary targeting of the heterotrimeric kinesin-2 is also RanGTP-dependent.

### RanGTP regulates ciliary length and ciliary trafficking of KAP under steady-state conditions in *Chlamydomonas*

To see if mechanisms of Ran-dependent ciliary targeting and entry are broadly conserved, we tested the effect of Ran manipulation on assembly and kinesin-2 motor targeting in *Chlamydomonas* cilia. The unicellular green algae *Chlamydomonas* is an excellent model to study ciliary length regulation and protein trafficking. In addition to the extensive body of literature on motor trafficking and ciliary assembly in this organism, the small G-protein Ran and key residues required for GTP hydrolysis are well conserved between humans and *Chlamydomonas* (**Figure S3**). We therefore examined the role of Ran-like protein (Ran1), the ortholog of human Ran, on ciliary length regulation in wild-type *Chlamydomonas* CC-125 cells. Importazole (IPZ), a small molecular inhibitor which specifically blocks RanGTP-importin β1 interaction (Soderholm et al.,2011), was used to perturb RanGTP function. The result indicated treatment CC-125 cells with IPZ for 2 hours shortens ciliary length in a dose-dependent manner (**Figure 5, A and B**).

**Figure 5.**
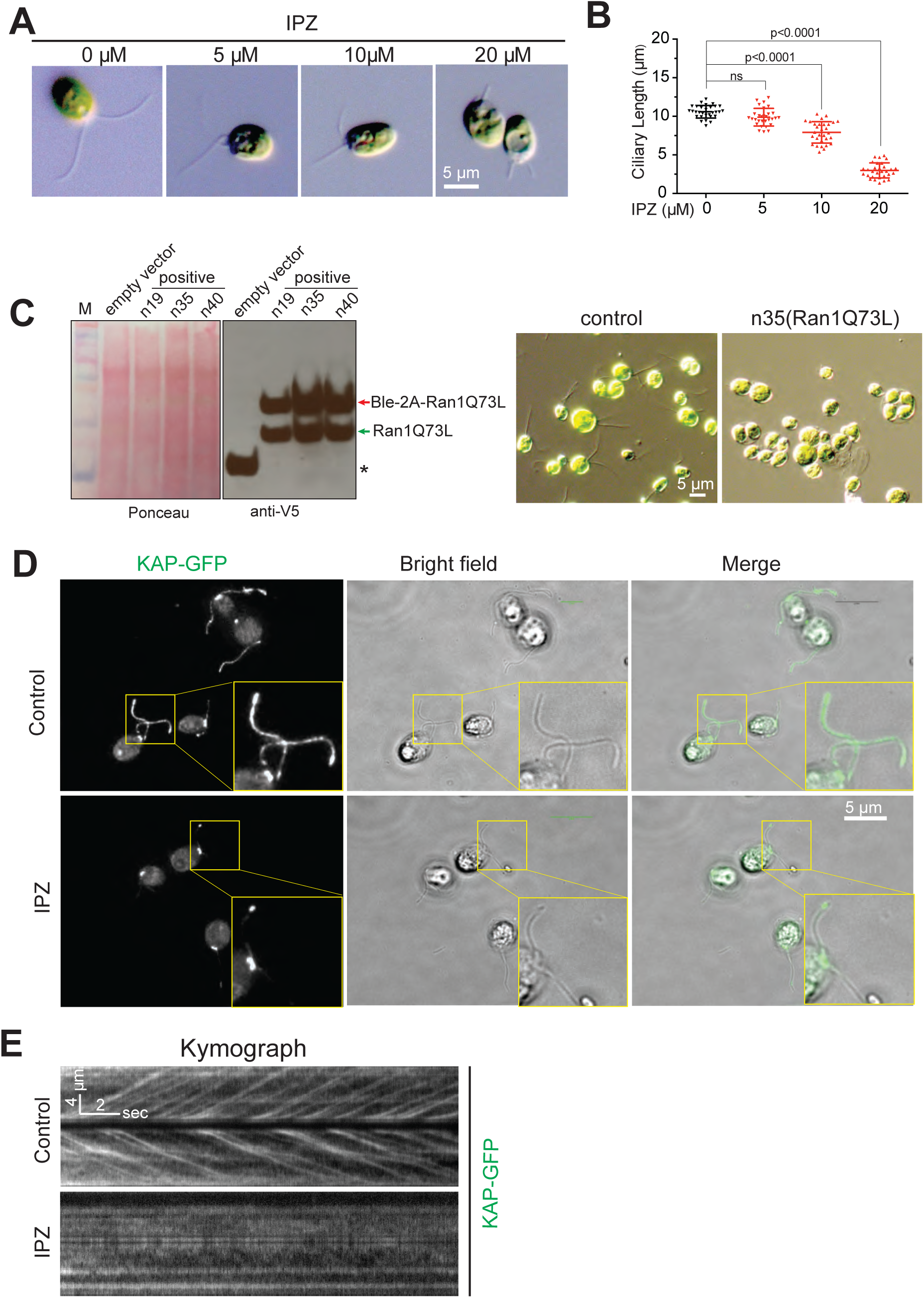
Inhibition of RanGTP function shortens cilia length and blocks ciliary trafficking of KAP under steady-state conditions in *Chlamydomonas*. **A**. Dose dependent inhibition of cilia length by the small molecular inhibitor IPZ, which disrupts RanGTP interacting with importin β. CC125 cells were treated with different concentration of IPZ for 120 min, fixed with 1% glutaraldehyde, and imaged by DIC microscope at 40× magnification. **B**. Quantification of cilia length. The unpaired *t*-test analysis was performed. *P*-value of less than 0.05 was considered significant difference. **C.** Expression of the dominant negative mutant Ran1Q73L, corresponding to human RanQ69L, exhibits either clumpy cells or shortened ciliary length in *Chlamydomonas.* Linearized expression plasmid pChlamy-4-Ran1Q73L was transformed into CC-125 cells and initially screened by colony PCR. The expression of Ran1Q73L was finally detected by western blotting analysis via V5 antibody. Due to cleavage efficiency of 2A peptide in *Chlamydomonas*, Ran1Q73L was existed in two forms: Ran1Q73L (correctly processed, green arrow) and Ble-2A-Ran1Q73L (fused with the selection marker, red arrow). Asterisk indicates that the negative control merely expresses zeocin resistance gene Ble. The representative images for the control Cells and Ran1Q73L expressing cells were shown. **D.** Inhibiting of Ran function by IPZ reduced ciliary localization of KAP. KAP-GFP reporter cells are treated with 20 μM IPZ for 2 h, fixed with 100% methanol, and mounted with prolong gold anti-fade mounting medium. Ciliary localization of KAP is dramatically reduced after IPZ treatment in *Chlamydomonas.* E. Kymograph analysis indicates IPZ blocked ciliary entry of KAP. KAP-GFP cells were treated with either DMSO or 50 μM IPZ for 20 min, and then used for TIRF microscopy to record the dynamic behavior of KAP.

To exclude that the phenotype was caused by the off-target effect of the inhibitor, the GTP-locked Ran1 mutant Ran1Q73L, corresponding to human RanQ69L, was transformed into *Chlamydomonas.* As shown in **Figure 5C**, there is strong Ran1Q73L expression as expected in *Chlamydomonas* (*green arrow*), despite a portion of expressed Ble-2A-Ran1Q73L fusion proteins being incompletely processed due to the cleavage efficiency of 2A peptide in *Chlamydomonas* (*red arrow*). Compared to control cells with normal ciliary length and cell division, the cells expressing high levels of Ran1Q73L exhibits either clumpy cells or shortened ciliary length. One possible explanation for clumpy cells may be that there are no cilia to secrete ectosomes containing lytic enzyme to break the cell wall after cell division (Wood et al., 2013). These results demonstrated that RanGTP plays pivotal roles in ciliary length regulation in *Chlamydomonas*.

We showed that KAP3, but not IFT81, couldn’t be targeted to ciliary base in hTERT-RPE cells constitutively expressing GTP-locked RanQ69L. It was reported that RanGTP regulates ciliary entry of the other motor KIF17, which is localized in the nucleus as KAP3 (Dishinger et al., 2010). Based on these data, we tested whether perturbing Ran function affected ciliary targeting or entry of KAP, the ortholog of human KAP3, in the *Chlamydomonas* KAP-GFP reporter strain CC-4296. As shown in **Figure 5D**, KAP-GFP is distributed in both the cilium and cilium base in control cells treated with DMSO. In contrast, ciliary localization of KAP-GFP was dramatically decreased in the cells treated with 20 μM IPZ for 2 hours. These results indicate that ciliary localization of KAP is regulated by RanGTP in *Chlamydomonas*. To further investigate whether IPZ directly affects ciliary entry of KAP, we used real-time total internal reflection fluorescence microscopy (TIRFM) to study the dynamic behavior of KAP. Kymograph analysis showed that KAP could enter into the cilium in un-treated cells. In contrast, ciliary entry of KAP is blocked in the cells treated with 50 μM IPZ (**Figure 5E**). The image shown was taken around 20 minutes after addition of IPZ indicating a direct role of RanGTP in regulation of ciliary entry of the motor subunit KAP as propagation of nuclear effects are likely to take longer than the immediate reduction in ciliary entry upon drug addition.

### RanGTP directly regulates ciliary protein incorporation during cilia regeneration in *Chlamydomonas*

One advantage of the *Chlamydomonas* model system in this context is the significant available information about requirements for nuclear regulation of ciliary assembly. During ciliary regeneration after ciliary severing (deciliation), new ciliary proteins need to be synthesized and transported to assembly sites for incorporation into cilia (**Figure 6A).** This process requires initiating gene expression, which would be dependent on nuclear import of specific transcription factors like XAP5 (Li et al., 2018). Therefore, it is possible that RanGTP regulates cilium length by indirectly affecting nuclear import and ultimately new transcription/ciliary protein synthesis.

**Figure 6.**
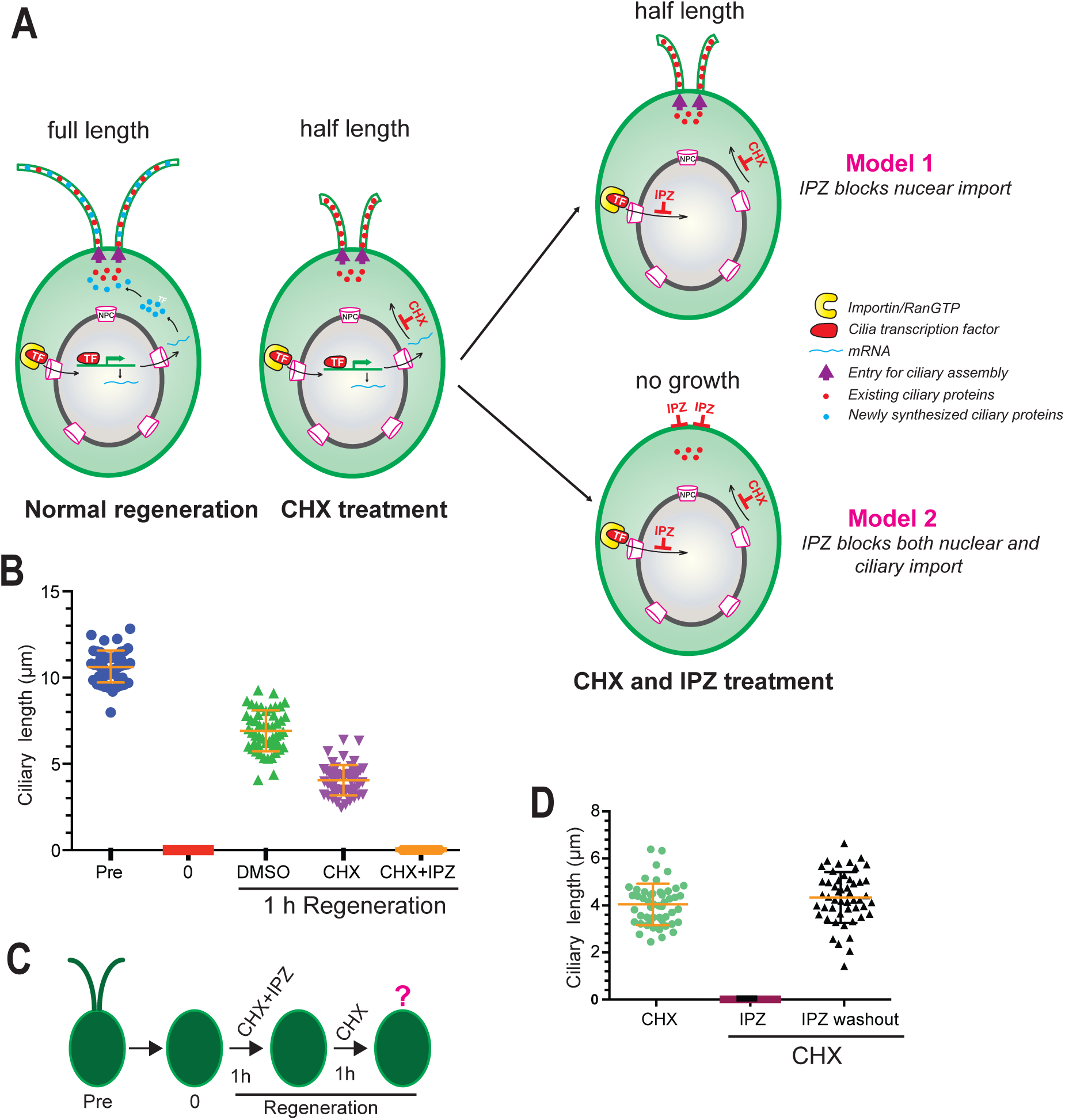
RanGTP directly regulates ciliary protein incorporation in *Chlamydomonas* during ciliary regeneration. **A.** Possible models of incorporation of existing ciliary protein into cilia after treated with cycloheximide (CHX)) and importazole (IPZ) during ciliary regeneration. Inhibition of new protein synthesis by CHX results in half-length cilia. **B.** Wild-type CC-125 cells were deciliated, and cilia were regenerated for 1 hour in the presence of different small molecular inhibitors: 10 μg/mL CHX, 10 μM IPZ or the combination of 10 μg/mL CHX and 10 μM IPZ. Cells were fixed with 1% glutaraldehyde and imaged by DIC microscope at 40×magnification. **C.** Schematic representation of IPZ washout assay under the absence of newly synthesized protein. **D.** Deciliated cells were treated with the combination of 10 μg/mL CHX and 10 μM IPZ for 60 min. Then IPZ, not CHX, was washed out, and cilia were regenerated for other 60 min. Cells were fixed with 1% glutaraldehyde and cilia length was measured using ImageJ software.

To tease apart nuclear and non-nuclear effects, we were able to use the small molecular inhibitor cycloheximide (CHX) to inhibit new protein synthesis during cilia regeneration. As shown in Figure 6A, in wild-type *Chlamydomonas* cells, this typically results in growth of cilia to half-length (6 µm) which exhibits the ability of these cells to incorporate already-synthesized proteins to generate half-length cilia without the production of new proteins from the burst of transcription post-deciliation (Rosenbaum et al., 1969). As expected, when blocking new protein synthesis with CHX, the existing ciliary proteins can build short cilia as shown in **Figure 6B**. If RanGTP exclusively inhibits nuclear import, but not ciliary import, inhibition of Ran function should allow existing ciliary proteins to still incorporate and assemble cilia to half-length (**Figure 6A**, **Model 1**). If inhibiting Ran function blocks both nuclear import and direct ciliary import, even the existing ciliary proteins shouldn’t incorporate and build cilia, resulting in bald cells (**Figure 6A, Model 2**). Our data fit Model 2 and show that when the deciliated cells are treated with IPZ to inhibit Ran function (with CHX to block any new protein synthesis), there is no cilium formation during regeneration. This demonstrates that IPZ can directly block incorporation of the existing ciliary proteins into cilia for assembly (**Figure 6B**). To confirm that the lack of ciliary growth wasn’t due to cell toxicity and that IPZ only impacts the ability of existing proteins to enter cilia, we washed out IPZ but still continued CHX treatment to inhibit new protein synthesis. As shown in **Figure 6 C and D**, ciliary biogenesis is restored upon IPZ washout. These data clearly showed that under conditions where only existing ciliary proteins can either enter cilia or not, RanGTP has direct effects in regulating ciliary protein incorporation. We also released CHX inhibition to confirm that, regardless of the presence of new proteins, blocking Ran function can regulate incorporation of existing ciliary proteins expected to enter cilia upon deciliation (**Figure S4 A and B**). Our data indicated IPZ can block incorporation of the existing ciliary proteins in addition to any newly synthesized proteins. In these conditions, if IPZ blocked nuclear entry of transcription factors needed for the spike in ciliary proteins but did not directly affect ciliary entry of existing proteins, cilia would still reach half-length from the already-synthesized ciliary protein pool. Ultimately, given the dual role of RanGTP in mediating ciliary import and nuclear import, it is important to segregate nuclear and direct ciliary effects of Ran perturbation. Here we are able to show that in spite of its demonstrated roles in regulating nuclear protein import, RanGTP has direct roles in mediating ciliary protein incorporation for cilia formation.

## Discussion

Although most kinesin motors are localized in the cytoplasm, different conditions allow some kinesin motors to transport into the nucleus including KAP3, KIF4, KIF17, and KIF17B (Morris et al., 2004; Seungoh et al., 2001; Dishinger et al., 2010; Macho et al., 2002). KAP3 and KIF4 can redistribute to the nucleus during mitosis (Morris et al., 2004; Seungoh et al., 2001). During mouse spermatid development, KIF17B shuttles from nucleus to cytoplasm (Macho et al., 2002). We observed that both the isoforms of the heterotrimeric kinesin-2 accessory subunit KAP3A and KAP3B are localized in the nucleus, and that their nuclear localization is RanGTP dependent. Considering KIF17B can function as a transcription regulator (Macho et al., 2002), it is possible that KAP3 participates in regulation of gene expression in the nucleus. Besides the kinesin motors, many cilia associate proteins can localize in the nucleus (Mcclure-Begley et al., 2017). The exchange of components between the ciliary and nuclear compartments is also thought to be mediated by membrane-less organelles (Johnson et al., 2019). However, the nuclear roles and origin of cilia associated proteins needs to be further investigated.

It was reported that the armadillo repeats of KAP3 are responsible for binding to motor subunits KIF3A/3B, and the C-terminal conserved domain is responsible for specific cargo binding (Haraguci et al., 2005; Nagata et al., 1998; Jimbo et al., 2002; Deacon et al, 2003). Our data showed the armadillo repeats 6-9 (ARM6-9) is required for KAP3 targeting to the ciliary base, probably mediated by RanGTP. It is possible that the heterodimeric KIF3A/3B and RanGTP collaboratively regulate KAP3 targeting to the ciliary base. It is also noteworthy that cells expressing the truncated KAP3A (186-660), containing only the armadillo repeat domain, have normal cilia length, whereas cells expressing the truncated KAP3 (186-792), containing both the armadillo repeats and cargo-binding domains, have no cilia. Given that loss of the cargo-binding domain dramatically decreases KAP3 binding to KIF3A/3B (Haraguci et al., 2005), our data suggest that the dominant negative function of KAP3 truncations is dependent upon their binding ability to the KIF3A/KIF3B motor subunits.

Several lines of evidence suggest that RanGTP is involved in ciliary protein trafficking (Dishinger et al., 2010; Hurd et al., 2001; Fan et al., 2011; Maiuri et al., 2013). RanGTP was reported to regulate ciliary entry of the homodimeric motor KIF17 and RP2 (Dishinger et al., 2010; Hurd et al., 2001). RanGTP was also reported to facilitate ciliary export of huntingtin Maiuri et al., 2013). However, there role of RanGTP on cilium formation was ambiguous. Two groups demonstrated that RanGTP has no effect on ciliary biogenesis (Dishinger et al., 2010; Torrado et al., 2016), whereas another group showed that manipulation of RanGTP concentration via RanBP1 knockdown could drive cilia formation (Fan et al., 2011). The difference of these observations for the role of RanGTP on cilia formation may be due to different cell lines, different strategies for inhibiting RanGTP function, or the timing and levels of RanGTP inhibition relative to cilia induction. Our results indicate that different dominant negative forms of Ran have different effects on cilia formation, although these mutants have no effect on regulating cilium length. Furthermore, GTP locked mutant RanQ69L more dramatically affects percent ciliation than that of RanG19V. The difference between RanQ69L and RanG19V is that RanQ69L has much higher affinity for GTP than RanG19V, thus RanQ69L-expressing cells have less free RanGTP than RanG19V-expressing cells. Taken together, our results suggest that precise manipulation of intracellular free RanGTP is critical for regulating cilium formation.

In addition to RanGTP, the importin transport receptors also participate in ciliary protein trafficking (Fan et al., 2007; Dishinger et al., 2010; Hurd et al., 2001; Torrado et al., 2016; Madugula et al., 2016; Han et al., 2017). There is also some disagreement about which importin is utilized for ciliary protein trafficking. Importin β1 was responsible for transmembrane protein Crumbs3 ciliary trafficking (Fan et al., 2007), and importin β2 was identified as the transport receptor for ciliary targeting of either transmembrane or soluble proteins like KIF17, Gli2 and GLi3 (Dishinger et al., 2010; Hurd et al., 2001; Madugula et al., 2016; Torrado et al., 2016; Han et al., 2017). However, additional data has shown that importin α1 and α6, but not importin β2, are responsible for ciliary targeting of soluble KIF17 (Funabashi et al., 2017). In general, importin β1, alone or in cooperation with importin α, transports substrates with a conventional NLS (Lange et al., 2007), whereas importin β2 transports substrates that contain the non-conventional PY-NLS (Lee et al., 2006). Consistent with this, ciliary targeting of the transcriptional factor Gli2/Gli3, which utilizes transport receptor importin β2, relies on its PY-NLS motif. PY-NLS mutations also result in the loss of Gli2/Gli3 ciliary targeting (Han et al., 2017). It is reported that the NLS-like sequence in the C-terminal region of KIF17 is required for its ciliary targeting (Dishinger et al., 2010). This NLS-like sequence was further confirmed as a classical mono-partite NLS (Funabashi et al., 2017). However, we noticed that PL, the PY variant, is located in the immediate downstream region of this NLS. So it is worth investigating whether or not this C-terminal NLS of KIF17 is a PY-NLS and which importin is used for KIF17 ciliary trafficking. Recently, a ternary complex consisting of importin β2, small GTPase Rab8 and ciliary targeting signals was reported to guide transmembrane protein trafficking to cilium (Madugula et al., 2016). This data suggests that spatial structure of the ternary complex, but not specific ciliary targeting sequences, are required for ciliary targeting of membrane proteins. This highlights that the detailed working model for how importin mediates ciliary import needs to be further clarified.

There is increasing data that nuclear import and ciliary import shares similar mechanisms, at least in part (Dishinger et al., 2010; Kee et al., 2012; Takao et al., 2014; Del Viso et al., 2016; Takao et al., 2017; Endicott et al., 2018). First, both the NPC and the ciliary pore complex form a diffusion barrier (Kee et al., 2012; Endicott et al., 2018). Second, the RanGTP/importin transport system is also used for ciliary protein trafficking (Fan et al., 2007; Dishinger et al., 2010; Hurd et al., 2011; Han et al., 2017). Third, some nucleoporins also localize in the ciliary base to regulate barrier diffusion ability (Dishinger et al., 2010; Del Viso et al., 2016; Endicott et al., 2018). Our data have shown that RanGTP can regulate cilium formation and ciliary trafficking of KAP3. One remaining critical question is whether RanGTP has direct effects in modulating ciliary protein transport. One possibility is that the effect of RanGTP is an indirect result of inhibiting nuclear import of proteins, like transcriptional factors, which are required for ciliary formation. By using the unicellular green alga *Chlamydomonas* as a model organism, we clearly demonstrated that RanGTP function directly regulates ciliary incorporation of the existing pool of already-synthesized ciliary proteins, which is not dependent on new transcription. In addition, the dominant negative mutant RanQ69L blocked ciliary trafficking of KAP3. Given that KAP is required for localization of KIF3A/3B to the assembly sites (Muller et al., 2005), RanGTP may control cilia formation by directly regulating ciliary targeting of the heterotrimeric kinesin-2 motor. This will in turn affect ciliary assembly and length maintenance due to the importance of ciliary recruitment and entry of kinesin-2 motor KIF3A/3B/KAP in these processes (Engel et al., 2009; Ludington et al., 2013). Further work will determine if this is a generalized mechanism for ciliary protein import and will identify additional RanGTP-regulated ciliary proteins (cargoes) required for cilium assembly, length control, and function.

## Materials and Methods

### Compounds

DMSO, Importazole (IPZ, #SML0341) and Cycloheximide (C1988) were purchased from Sigma-Aldrich. Indicated concentrations and specific incubation times are used in this study.

### DNA constructs

Plasmids for HA-tagged human KAP3A and KAP3B were kindly from Dr. Benjamin Allen (University of Michigan). Plasmids for wild-type Ran and point mutants RanG19V and RanT24N are a generous gift from Dr. Kristen Verhey (University of Michigan). Plasmids expressing MBP and M9M were from Dr. Yuh Min Chook (University of Texas Southwestern Medical Center). Plasmids pmCherry-C1-RanQ69L (#30309) was obtained from Addgene under the material transfer agreement. GeneArt^TM^ *Chlamydomonas* protein expression vector pChlamy_4 was from Therm Fisher Scientific. Recombinant plasmids pChlamy_4_Ran1Q73L, EGFP or HA-tagged KAP3 truncations were generation by ligation-independent cloning strategy as described before (Zhu et al., 2010) and sequenced in full.

### *Chlamydomonas* strains, mammalian cells and antibodies

Wild-type and KAP-GFP reporter strains were obtained from the *Chlamydomonas* resource center (CC-125 mt^+^ and CC-4296). Strains were grown in liquid Tris-Acetate-Phosphate (TAP) liquid medium for 18-24 hours prior to experimentation. Mammalian COS-7 and MDCK Cells were cultured in Dulbecco’s Modified Eagle’s Medium (DMEM; Invitrogen) supplemented with 10% fetal bovine serum (FBS; Invitrogen). Human TERT-RPE cells were cultured in DMEM+F12 (1:1) (Invitrogen) containing 10% FBS. Antibodies used in this study are as follows (IF and WB are short for immunofluorescence and western blot, respectively): Mouse anti-acetylated α-tubulin (#T6793, 1:500 for IF) was from Sigma-Aldrich (St. Louis, MO, USA). Rabbit anti-Cep164 (#22227-1-AP, 1:50 for IF) and rabbit anti-IFT81 (#11744-1-AP, 1:50 for IF) were from Proteintech. Mouse anti-KAP3A (#610637, 1:20 for IF) was from BD Transduction Laboratories™. Mouse anti-Myc (AB_390912, 1:100 for IF) was from Roche. Mouse anti-hnRNP A1 Antibody (sc-32301, 1:50 for IF) was from Santa Cruz biotechnology. Rabbit anti-myc (#5625, 1:500 for IF), Rabbit anti-V5 (#13202, 1:1000 for WB), rabbit anti-HA (#3724, 1:100 for IF) and Rabbit anti-GFP (#2956, 1:100 for IF and 1:1000 for WB, respectively) were from Cell Signaling Technology.

### Cell culture and transfection

COS-7, MDCK and hTERT-RPE cells were maintained in a humidified atmosphere at 37°C and 5% CO2. Cells for transfection were seeded in an 8-well chamber slide (Lab-Tek) with 0.4 mL culture medium per well. After overnight growth, the cells became 70-80% confluent and were transfected with the corresponding plasmids using the transfection reagent FuGENE 6 (Roche) according to the manufacturer’s instructions. In normal condition, COS-7, MDCK and hTERT-RPE cells are fixed with 4% paraformaldehyde for intracellular localization assay 24 hours post-transfection. In serum starvation condition for cilium induction, hTERT-RPE cells were cultured in complete medium for 24 hours post-transfection, then followed to culture in DMEM+F12 (1:1) with 0.25% FBS for other 24 hours.

### *Chlamydomonas* transformation

Electroporation transformation was used form rapid transformation of *Chlamydomonas* with the electroporator NEPA (Nepa Gene, Japan). Transformation was performed following the published protocol with some modifications (Yamano et al., 2013). The typical 4 days were necessary to perform the transformation. *Day 1*: pre-cultivation stage: the cells were grown in 5 mL TAP liquid medium for overnight culture. *Day 2*: pre-cultured cells from day 1 were transferred into a new 50 mL TAP medium in a 250 mL flask with a final OD_730_ of 0.1 (usually 1-3 mL pre-cultures added) for overnight culture with 120 rpm/25°C. *Day 3*: cells were harvested by centrifugation when the cell density reached OD_730_ of 0.3-0.4, and washed by GeneArt MAX Efficiency Transformation Reagent (Invitrogen) 3 times and re-suspended in 250 μL TAP medium containing 40 mM sucrose. 1.6 μg linearized DNA (pChlmay_4_Ran1Q73L) was mixed with 160 μL of the cell suspension for electroporation. After electroporation, the cells were transferred into 10 mL TAP plus 40 mM sucrose for overnight culture in dim light. *Day 4*: cells were collected and plated onto 1.5% TAP-agar plate with 10 μg/mL zeocin for growth. The colonies will be visible 5-7 days later.

### Immunofluorescence staining

Cells were washed with cold PBS twice, and then fixed with 4% paraformaldehyde in HEPES (pH 7.4) for 15 min at room temperature. Cells were washed three times with cold PBS and then incubated with 0.1% Triton X-100 in PBS (pH 7.4) for 10 min. Permeabilized cells were washed with PBS three times, and then incubated in PBS with 10% normal goat serum and 1% BSA for 1 hour at room temperature to block non-specific binding of the antibodies. Cells are incubated with diluted primary antibody in PBS with 1% BSA overnight at 4°C. After three times wash with PBS, cells are incubated with the secondary antibody in PBS with 1% BSA for 1 hour at room temperature in the dark. After washing three times with PBS, cells are mounted with ProLong Antifade mounting medium with or without DAPI, and kept at 4°C in the dark for further imaging.

### TIRF Microscopy

Samples were prepared as follows: KAP-GFP reporter cells (CC-4296) were cultured in TAP liquid medium for 18 h, then centrifuged at 1000 rpm for 2 min. 3 μL cell pellets were re-suspended in 200 uL TAP liquid medium containing either DMSO or 50 uM IPZ. 24 × 50 mm no. 1.5 coverslip was treated with 0.1% poly-lysine for 10 min, and dipped into water and let it air-dry. Petroleum jelly was used to draw a circle around poly-lysine coated region. 20 μL of cells were placed inside and allowed to settle for 3 min, and then imaged. KAP-GFP cells were imaged on a Nikon Ti-E microscope with a 100x 1.49 NA oil objective. Images were obtained at 23.95 fps with 0.16 μm per pixel by using an Andor DU897 electron multiplying charge-coupled device (EMCCD) camera. The NIS-Elements software was used to crop the movies and generate kymographs.

### Ciliary regeneration

*Chlamydomonas* cells were deciliated by pH shock as described before (Witman et al., 1972), and ciliary regeneration was induced in normal TAP liquid medium. After deciliation, cells were immediately treated with 10 μg/mL Cycloheximide and/or 10 μM IPZ for 60 min. For IPZ washout experiment, treated cells were washed 3 times and cultured in fresh TAP liquid medium (or with 10 μg/mL Cycloheximide). Cells were fixed with 1% glutaraldehyde for 15 min at room temperature, and cilia length was measured using the line segment tool in ImageJ.

### Statistical analyses

All data are reported as mean values ± standard error of the mean (S.E.M). Graphs and associated statistical analyses were performed with Prism 6.0C (GraphPad Software, La Jolla, CA). The unpaired student’s t-test was used to assess statistical significance of two groups. A value of p < 0.05 was considered statistically significant.

## Acknowledgements

We would like to thank Dr. Yuh Min Chook (University of Texas Southwestern Medical Center), Drs. Kristen Verhey and Benjamin Allen (University of Michigan) for providing DNA plasmids. We also thank our colleague Dr. Pamela Tran for sharing some reagents. We are grateful to the members of Dr. Avasthi Lab for comments on the manuscript and Dr. Pawel Niewiadomski (University of Warsaw) for feedback on our preprint. We would like to thank Ms. Larissa Dougherty for editing the manuscript. This work was supported by the following grants: startup of Department of Ophthalmology (P.A.) and the Biomedical Research Training Program (S.H.), University of Kansas Medical Center; a postdoctoral fellowship from a Kansas Institutional Development Award (IDeA) from the National Institute of General Medical Sciences of the National Institutes of Health under grant number P20 GM103418 (S.H.).

---

**Figure S1.**
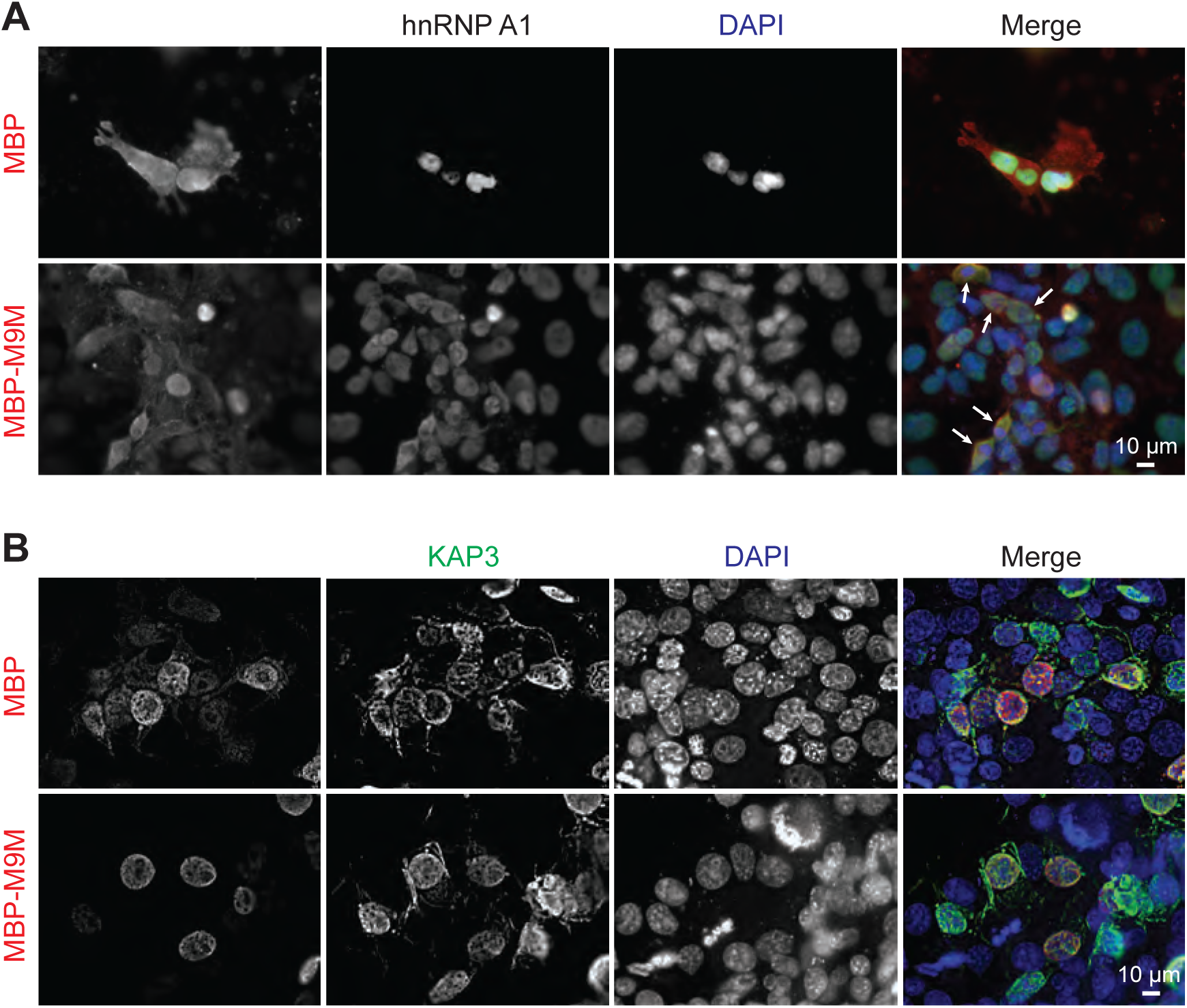
Nuclear translocation of KAP3 is transport receptor importin β2 independent. **A.** Importin β2 mediates nuclear translocation of hnRNP A1. The plasmid myc-MBP-M9M expressing myc-MBP fused inhibitory peptide M9M of importin β2, or its control merely expressing myc-MBP, was transfected into COS-7 cells. 24 hours after transfection, cells were fixed with 4% PFA and co-stained with rabbit anti-myc (Red) and mouse anti-hnRNP A1 antibodies (green). nuclei are stained with DAPI (blue). **B.** Importin β2 isn’t required for nuclear localization of KAP3. Importin β2 mediates nuclear translocation of hnRNP A1. The plasmid myc-MBP-M9M or its control was co-transfected with EGFP-KAP3 into COS-7 cells. 24 hours after transfection, cells were fixed with 4% PFA and co-stained with mouse anti-myc (Red) and rabbit anti-EGFP antibodies (green). nuclei are stained with DAPI (blue).

**Figure S2.**
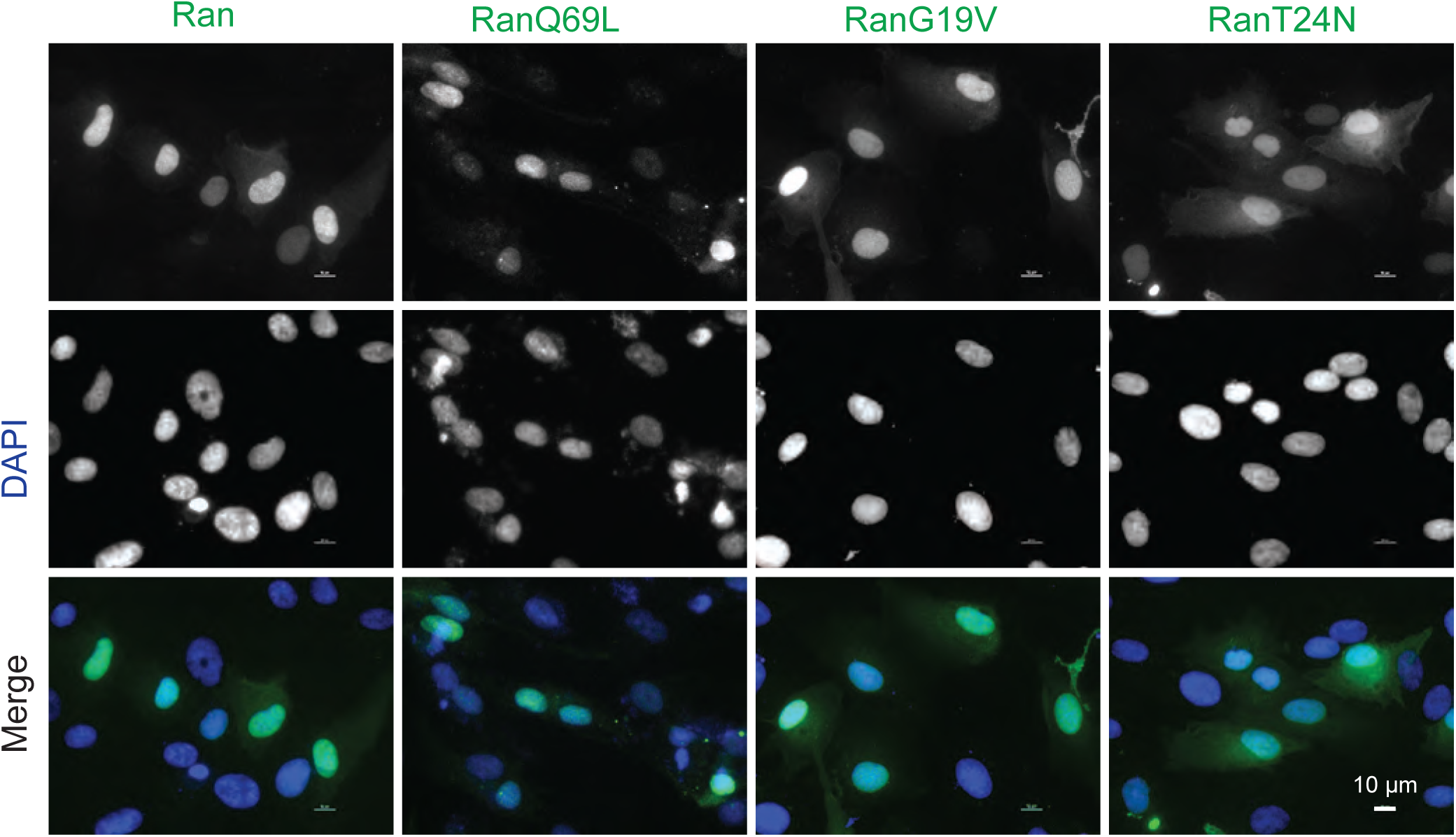
Subcellular localization of wild-type Ran and its dominant negative mutants in serum starved hTERT-RPE cells. Plasmids for expressing wild-type Ran or its point mutants (RanQ69L, RanG19V and RanT24N) were transfected into hTERT-RPE cells. After 24 h transfection, the cells were suffered for serum starvation for another 24 h, and then fixed with 4% PFA. The intracellular localization of these proteins was visualized via immunofluorescence staining.

**Figure S3.**
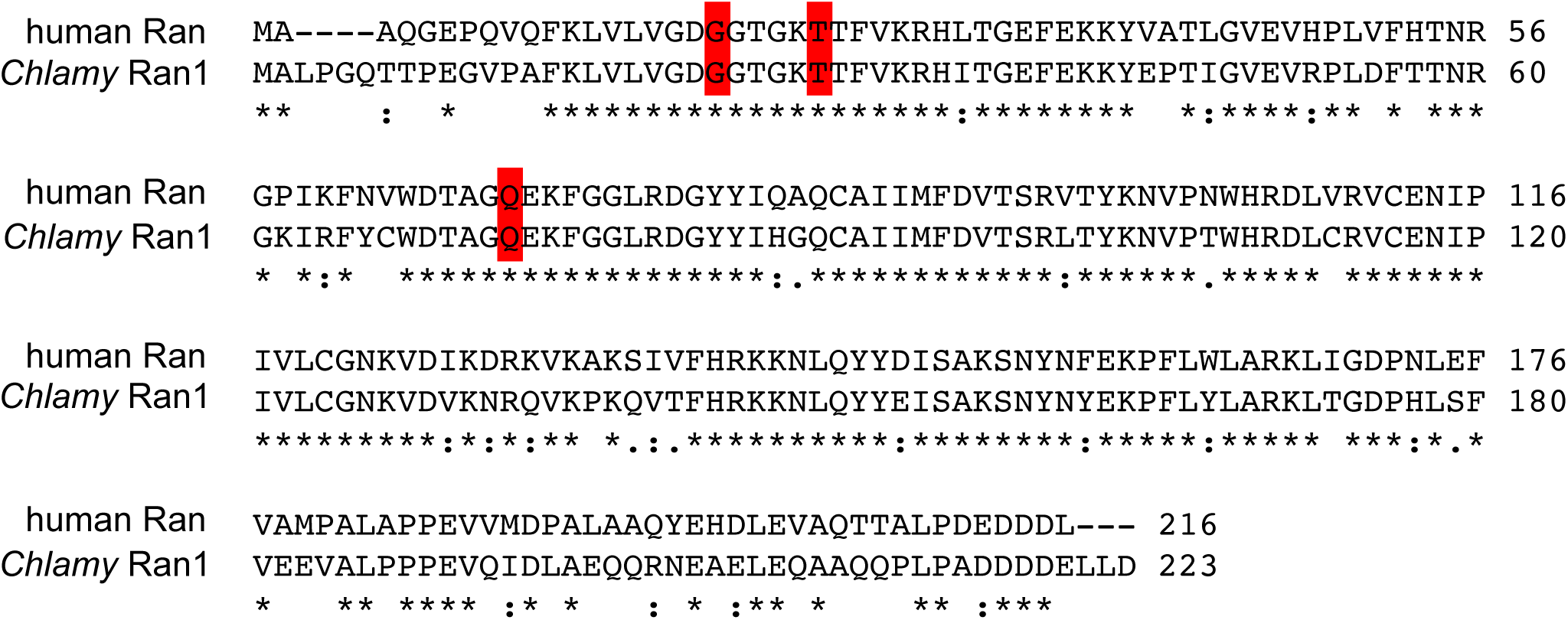
Alignment of human Ran with *Chlamydomonas* Ran like small GTPase (Ran1). Human Ran (NCBI reference sequence: NP_006316.1) and *Chlamydomonas* Ran1 (Phytozome reference sequence: cre03.g191050.t1.2) were aligned using the Clustal Omega software. Conserved residues are indicated (asterisk indicates fully conserved residues; colon indicates residues with strongly similar properties; period indicates residues with weakly similar properties). The key residues required for GTP or GDP bound state of Ran are marked in red.

**Figure S4.**
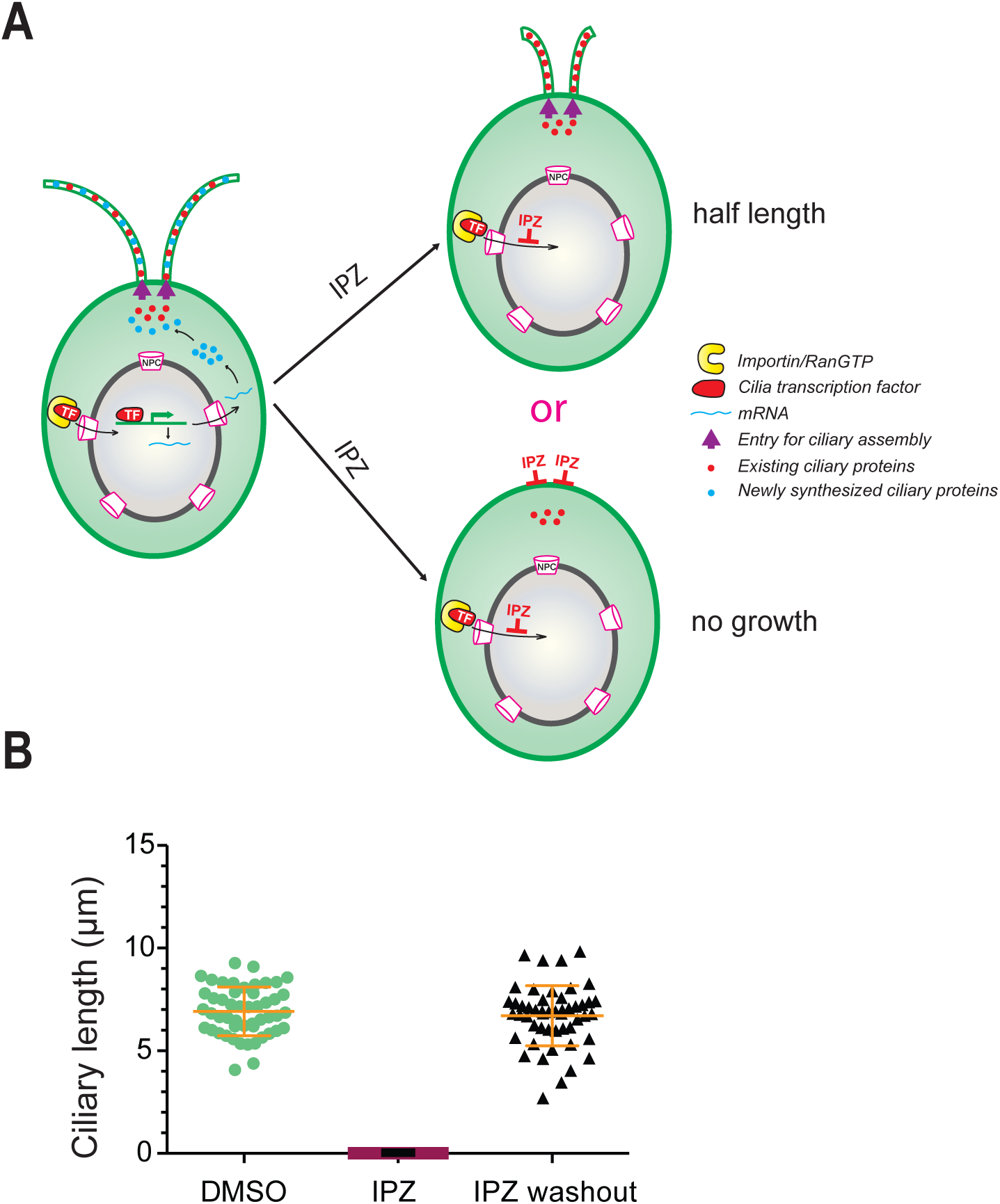
RanGTP regulates ciliary protein incorporation in *Chlamydomonas* regardless of the presence of new synthesized proteins. **A.** Possible models for RanGTP regulating ciliary protein incorporation in the presence of new synthesized proteins. **B.** Deciliated cells were treated 10 μM IPZ for 60 min. Then IPZ was washed and cilia were regenerated for another 60 min. Cells were fixed with 1% glutaraldehyde and cilia length was measured using ImageJ software.

